# Molecular evidence indicates the existence of multiple lineages of *Sperata species* in Indian Rivers

**DOI:** 10.1101/2020.11.16.377507

**Authors:** Prabhaker Yadav, Ajit Kumar, Neha Yadav, Mansi Bisht, Syed Ainul Hussain, Sandeep Kumar Gupta

## Abstract

*Sperata seenghala* (Giant river-catfish) and *Sperata aor* (Long-whiskered catfish) are commercially important freshwater catfishes of India, belongs to family Bagridae. Due to high nutritional significance and the low number of intramuscular bones, both fishes have considerable demand in South Asian countries. Both of the *Sperata* species are morphologically close and well adapted to the same habitat. In this study, we have assessed the level of genetic diversity and differentiation of *S. seenghala* and *S. aor* in the Ganga River based on the mitochondrial DNA (mtDNA) control region and compared with the other major Indian rivers. We found high haplotypes diversity for both the species in the Ganga. However, it was comparatively low for *S. seenghala* in Mahanadi and Brahmaputra populations. The phylogenetic and median-joining network strongly indicated the presence of two distinct maternal lineages of *S. seenghala* from the Ganga river. Interestingly, the genetic differentiation between *S. seenghala* of Ganga-Brahmaputra was much higher (~25.3%) than the *S. seenghala* and *S. aor* (~17%), whereas it was comparatively low between Ganges-Mahanadi (~8.0%). Our finding provided evidence that all the three rivers: Ganga, Mahanadi, and the Brahmaputra sustain a highly diverse and genetically distinct stock of giant river catfish; therefore, all populations should be considered as a different management unit for the protection of stocks. Our findings indicated that Brahmaputra lineages qualify the species level variations. This study can be further used as a reference database for proper lineage identification of *S. seenghala* and *S. aor* that could formulate the appropriate conservation and management plans.

## Introduction

The genus *Sperata* is one of the largest catfish that belongs to family Bagridae and widely distributed in India, Pakistan, Bangladesh, Afghanistan, and Nepal (Talwar & Jhingharan, 1991). The genus *Sperata* was previously known as *Aorichthys*/*Mystus* and was recognized as one of the commercially important catfish species (Saigal & Motwani, 1961; Froese & Pauly, 2011). Due to high nutritional values and the low number of intramuscular bones, *Sperata* species have a huge demand in South Asian countries (Mohanty *et al.*, 2012). Based on the morphological characteristics, four species of *Sperata* are currently recognized from Asia (Day, 1878; Talwar & Jhingharan, 1991; Ferraris & Runge, 1999; Ng & Kottelat, 2013). These four species are named as, *S. seenghala* (Sykes, 1839), *S. aor* (Hamilton, 1822), *S. acicularis* (Ferraris & Runge, 1999), and *S. aorella* (Blyth, 1858). Recently, there are one more species, *S. sarwari,* which is recognized from the Indus river system of India and Pakistan (Nawaz *et al.*, 1994). The catfish can tolerate a high range of temperature (38-40 °C), low dissolved oxygen, and salinity along with variable water conditions, because of their robust air-breathing system (Yadav *et al.*, 2017; 2018). Among the four species of the genus *Sperata*, *S. seenghala* (the giant river-catfish) and *S. aor* (the long-whiskered catfish) have a wide distribution in the Indian subcontinent (Gupta, 2015). In the Ganga river, due to the high anthropogenic activities such as extensive fishing, water abstraction, pollution, siltation, and invasion of exotic species, are threatening the *Sperata* populations. *S. seenghala* and *S. aor* are recognized as “least concern” in the RedList of International Union for Conservation of Nature (IUCN). Previous studies on giant river catfish gradually focused on the genetic assessment of *S. seenghala* using Randomly amplified polymorphic DNA (RAPD) marker (Saini *et al.*, 2010; Garg *et al.*, 2014), mtDNA markers (Kumari *et al.*, 2017), and microsatellite marker (Acharya *et al.*, 2019). The microsatellite-based stock discrimination of *S. aor* revealed the existence of three genetic stocks from the connected vicinity of the Ganga river (Nazir & Khan, 2017). Though both the species of *Spereta* are abundantly found in Indian rivers; no study has been done so far to differentiate them.

Day (1878) documented the morphological characteristics for the identification of these catfish species. In *S. aor* maxillary barbels are longer in size and prolonged or beyond to the base of the caudal fin. It poses a supraoccipital spine, and its interneural shield shares almost the same length. It is also characterized by 10-11 pectoral-fin rays, 19-20 gill rakers, and its orbit prolongs through the middle of the length of the head. However, in *S. seenghala*, the snout is chisel-shaped; maxillary barbels are not extended beyond the middle body; and supraoccipital spine shorter than interneural shield. It possess 8-9 pectoral-fin rays, 13-15 gill rakers, and its orbit completely present in the head’s anterior half (Gupta, 2015; Miyan *et al.*, 2016). Despite being significant variations, the identification between these two species is quite difficult. Hence, reliable sampling is essential for resolving the population genetic structure and diversity. Therefore, an appropriate management plan for a different stock can be prepared by incorporating reliable knowledge about existing lineages from species distribution ranges. Due to the maternal mode of transmission, and more rapid evolution than the nuclear genome, mitochondrial DNA (mtDNA) provides reliable information on the relationships among closely related species and populations (Brown *et al.*, 1979). In particular, the mtDNA control region (CR) is a highly variable portion and estimated to be five times more substitution rate than that of the rest sequences in mtDNA (Aquadro & Greenburg, 1983). For this reason, this marker is widely acceptable for population genetics studies (Wan *et al.*, 2004; Gupta *et al.*, 2018).

We used *S. seenghala* and *S. oar* from the Ganga river to improve our knowledge of contemporary genetic relationships, diversity, and demography of *Sperata*. Then we compared it with *S. seenghala* populations of Mahanadi and Brahmaputra. We used across geographic sampling from different locations of the Ganga River and DNA sequences of other major rivers for comprehensive genetic insight into *Spereta* lineages.

## Materials and methods

A small portion of fin tissues was collected from *S. seenghala* (n=131) from eight locations and *S. aor* (n=78) from four locations of the Ganges river and stored in 70% ethanol at 4°C. (Table 1, Fig. 1). The biological sample was collected from local fishermen along the river being harvested for selling in the fish market, and no animals were captured explicitly for this study. Therefore, Institutional Animal Ethics approval was not required for this research. Total genomic DNA was extracted from the tissue samples using the phenol-chloroform extraction protocols with a final elution volume of 100 μl (Sambrook *et al.*, 1989). The extracted DNA was checked on 0.8% agarose gel and quantified in QIAxcel and diluted in a final concentration of 30ng/μl for PCR amplification.

**Table 1.**
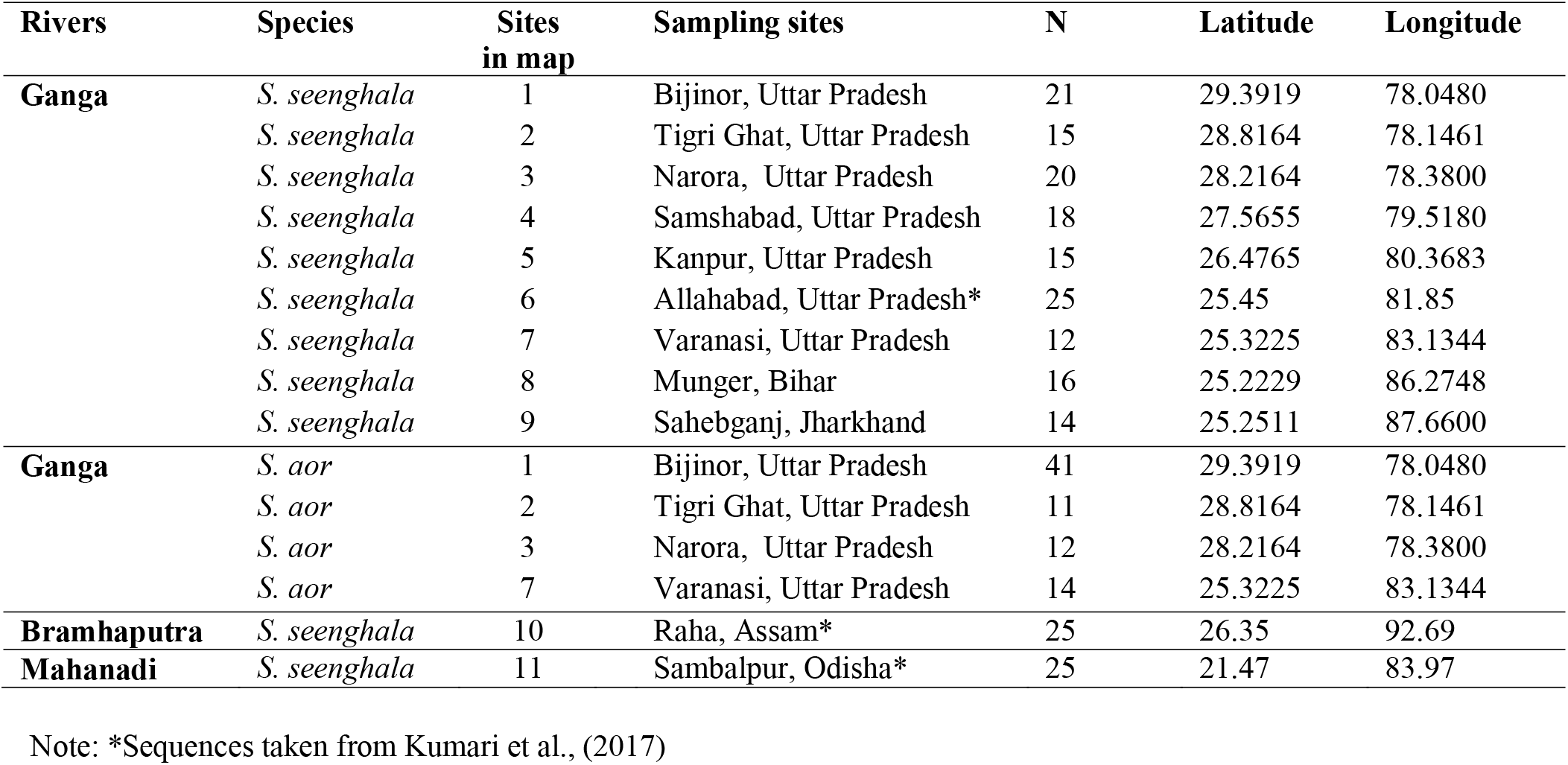
Sampling locations of *S. seenghala* and *S. aor* used in this study. N, number of samples.

**Figure 1.**
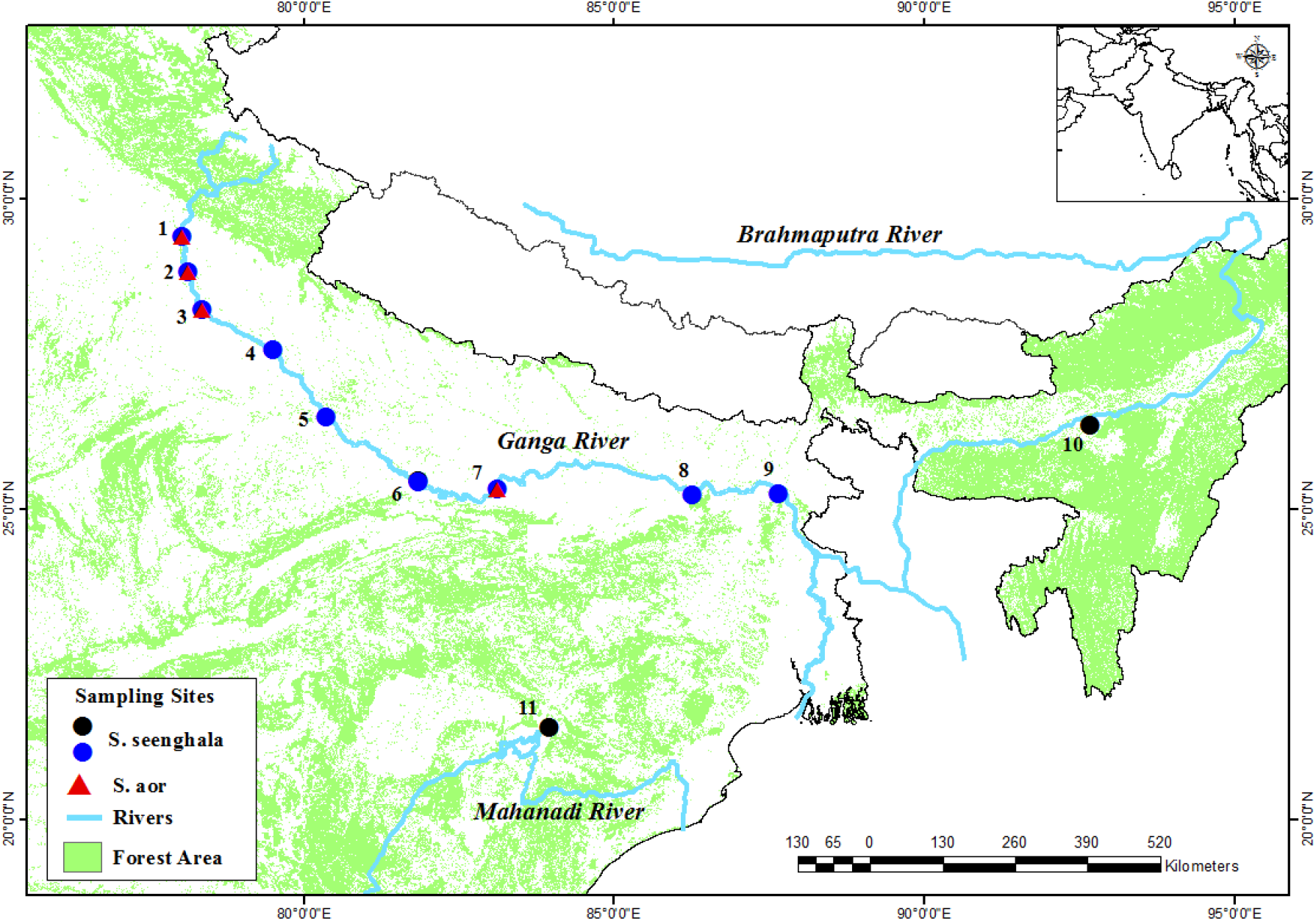
The Ganga, Brahmaputra and Mahanadi river system in India and the geographic origin of *Spereta seenghala* and *S. aor* samples used in this study.

### PCR amplification and sequencing

PCR amplification was performed using mtDNA control region (CR) primers SSDLoop F:5◻-CACCCCTAACTCCCAAAGC-3◻, and SSDloop R:5◻-GGTTTAGGGGTTTAACAGG-3◻(Kumari *et al.*, 2017). PCR reaction was performed in total reaction volumes of 20 μl using a 1 X PCR buffer (10 mM Tri –HCl, pH 8.3, and 50 mM KCl), 1.5 mM MgCl_2_, 0.2 mM of each dNTP, 2 pmol of each primer, 5 U of Taq DNA polymerase and 1 μl (~30 ng) of the template DNA. The PCR conditions were as follows: 95°C for 5 minutes, followed by 35 cycles of denaturation at 95°C for 45 seconds, annealing at 58°C for 45 seconds, and extension at 72°C for 75 seconds. The final extension was at 72°C for 10 minutes. The consistency of the PCR amplification was monitored using positive and negative controls. The PCR amplicons were checked on 2% agarose gel by gel electrophoresis and visualized under the UV transilluminator. Exonuclease I (EXO-*I*) and shrimp alkaline phosphatase (SAP) treatments were given to the amplified PCR products (USB, Cleveland, OH) for 15 min at 37°C and 20 min at 80°C, to degrade any residual primer and dNTPs. The amplified PCR products were sequenced from both forward and reverse directions using the BigDye® Terminator kit (v3.1) and analyzed on an Applied Biosystems Genetic Analyzer ABI 3500XL.

### Data analysis

All the raw sequences obtained from the forward and reverse directions were aligned and edited with SEQUENCHER®version 4.9 (Gene Codes Corporation, Ann Arbor, MI, USA) to get the consensus sequence. All the consensus sequences were aligned using the program CLUSTAL X multiple sequence alignment (Thompson *et al.*, 1997) and examined by visual inspection. DnaSP 5.0 (Librado & Rozas, 2009) was used to analyze the haplotype diversity (h), nucleotide diversity (p), and polymorphic sites (s). The numbers of nucleotide substitutions per site were estimated for multiple substitutions using the Tamura-3 parameter method in MEGA X (Kumar *et al.*, 2018). For the genetic distance, we used the Tamura-3 parameter using a discrete Gamma distribution (TN92+G) with the lowest BIC score value using MEGA X (Kumar *et al.*, 2018). Phylogenetic analyses were conducted in BEAST ver 1.7 (Drummond *et al.*, 2012). The spatial distribution of haplotypes was visualized by a median-joining network, was created with the PopART software (Leigh & Bryant, 2015). MEGA X (Kumar *et al.*, 2018) was used to estimate within-group genetic distance and between-group mean distance between the populations of *S. seenghala* and *S. aor*. In addition, molecular variance analysis (AMOVA) was performed to test the genetic differentiation between geographical units. The significance values generated by AMOVA were tested by random permutations of sequences among populations. DnaSP 5.0 (Librado & Rozas, 2009) was used to generate the mismatch distribution plot for trends in spatial demography history of *S. seenghala* and *S.aor* populations. Besides, to determine the demography history of each population of both the species, we have performed neutrality test, Tajima’s D (Tajima, 1989), Fu’s Fs test (Fu, 1997), the sum of squared deviations (SSD) and Raggedness index (r) under the demographic expansion model for each population using the program Arlequin 3.5 (Excoffier & Lischer, 2010). The P-values were obtained by 1000 simulations based on a selective neutrality test.

## Results

### mtDNA control region sequence polymorphism and haplotype diversity

We obtained 870 bp sequences for CR from 131 samples of *S. seenghala* from eight different sites and 78 samples of *S. aor* from four sites of the Ganga river (Table 1, Figure 1). Besides, 25 sequences of *S. seenghala,* each from the Ganga river, Mahanadi river, and Brahmaputra river were included from NCBI GenBank (Table 1, Supplementary Table: ST 1). Therefore, a dataset of 156 sequences of *S. seenghala* from the Ganga river were used in the present analysis. The high number of segregating sites was found in the Brahmaputra (173) followed by Mahanadi (136), *S. seenghala* of Ganga (86), and *S. aor* of Ganga (42) population. When sequences of all three rivers were aligned with our data, all were grouped into 77 haplotypes (Supplementary Table ST1). Out of these, 37 haplotypes (Hap1 to Hap 37) were observed in *S. seenghala* from the Ganga, 28 haplotypes (Hap38 to Hap65) were found in *S. aor*, four haplotypes (Hap 66 to Hap 69) were observed in Mahanadi, nine haplotypes (Hap3, Hap70 to Hap77) were observed in Brahmaputra river. Interestingly, four haplotype (Hap34 to Hap37) from the Ganga river sustain distinct genetic signature and highly diverse than the sequences of *S. seenghala* from the Ganga. The newly identified haplotypes of *S. seenghala* and *S. aor* from the Ganga River were submitted in GenBank (MTXXXXXX- MTXXXXX).

The haplotype diversity (hd) of *S. seenghala* and *S.aor* from the Ganges river were comparable and high 0.943 and 0.934, respectively. In contrast, the haplotype diversity of the Mahanadi and Brahmaputra rivers was 0.717 and 0.820, respectively. The nucleotide diversity (π) was ranging from 0.009 in *S.aor* to 0.049 in the Mahanadi population. The ratio of transition and transversion rate (Ts/Tv) was high in Mahanadi (94/49), whereas it was low in Brahmaputra 11/15 (Table 2).

**Table 2.** Haplotype diversity of *S. seenghala* and *S. aor* within rivers.

| Parameters | *Sperata seenghala* |  |  | *Sperata aor* |
| --- | --- | --- | --- | --- |
|  | Ganga | Mahanadi | Brahmaputra | Ganga |
| **n** | 158 | 25 | 25 | 78 |
| **N<sub>h</sub>** | 37 | 4 | 9 | 28 |
| **S** | 86 | 136 | 173 | 42 |
| **Ts/Tv** | 69/19 | 94/49 | 11/15 | 37/5 |
| **h (±SD)** | 0.943 (0.008) | 0.717(0.056) | 0.820 (0.065) | 0.934(0.016) |
| **π (±SD)** | 0.012 (0.001) | 0.049(0.012) | 0.034(0.017) | 0.009(0.0008) |
| **Neutrality Analysis** |  |  |  |  |
| **Tajima's D (P)** | -1.314(0.061) | 0.498(0.669) | -1.149(1.02) | -0.295(0.399) |
| **Fu's Fs (P)** | -4.662(0.152) | 31.053(1.00) | 1.107(0.717) | -4.900(0.109) |
| **Mismatch analysis** |  |  |  |  |
| **SSD (P)** | 0.2500(0.770) | 0.171(0.00) | 0.492(0.60) | 0.014(0.480) |
| **r-index (P)** | 0.0062(0.900) | 0.261(0.00) | 0.058(1.00) | 0.007(0.910) |

Note: Sample size (n); number of haplotypes (N<sub>h</sub>); segregating sites (S); haplotype diversity (h); nucleotide diversity (π); Standard diversity (SD); sum of square deviation (SSD); raggedness index (r)

### Molecular phylogeny and network

The Bayesian consensus tree was generated to access the genetic relationships between all haplotypes belongs to Ganga, Mahanadi, and Brahmaputra (Fig. 2). The phylogenetic tree showed that *S. seenghala* from the Ganga formed two distinct clades: clade-I A and clade-I B, whereas Mahanadi forms a basal clade II. The *S. aor* and the Brahmaputra sequences formed distinct clade-III and IV. Two sequences of Brahmaputra (Hap 3) were merged with the haplotype of Ganga *seenghala,* and three sequences of Mahanadi (Hap 69) were clustered with haplotype of *S. aor* (Supplementary Table ST1 and Fig 2). The median-joining network of all recognized haplotypes strongly indicated the presence of multiple lineages in *S*. *seenghala* from the studied river (Fig. 3). The clustering pattern obtained from network analysis is highly supportive of phylogenetic analysis.

**Figure 2.**
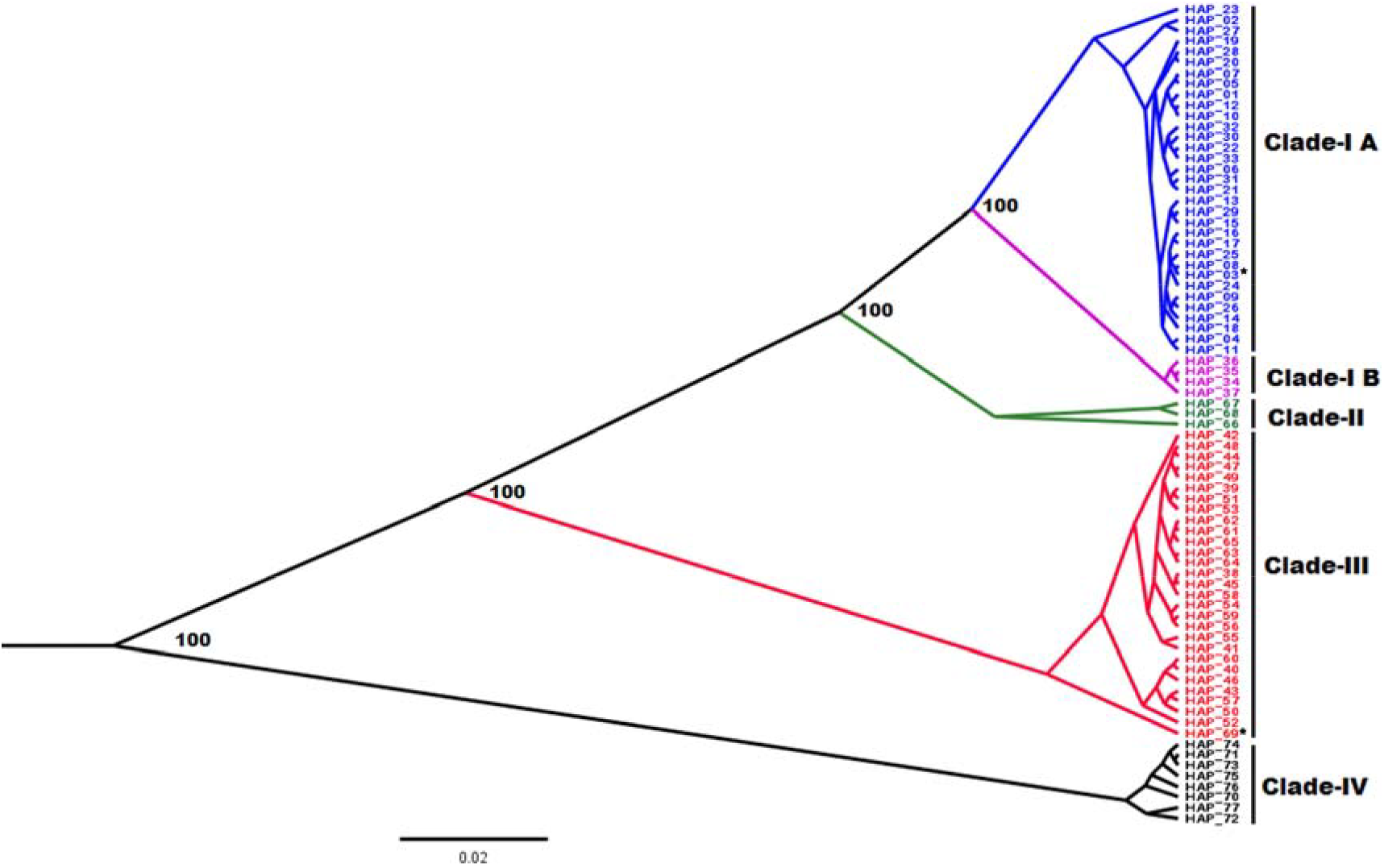
The Bayesian tree for the phylogenetic resolution between the *S. seenghala* form Ganges, Mahanadi, Brahmaputra, and *S. aor* Ganges. Numeric value at each node represents the posterior probability. Asterisk (*) represents, Hap3 belongs to Brahmaputra and Hap69 belongs to Mahanadi population identified by Kumari et al., (2016).

**Figure 3.**
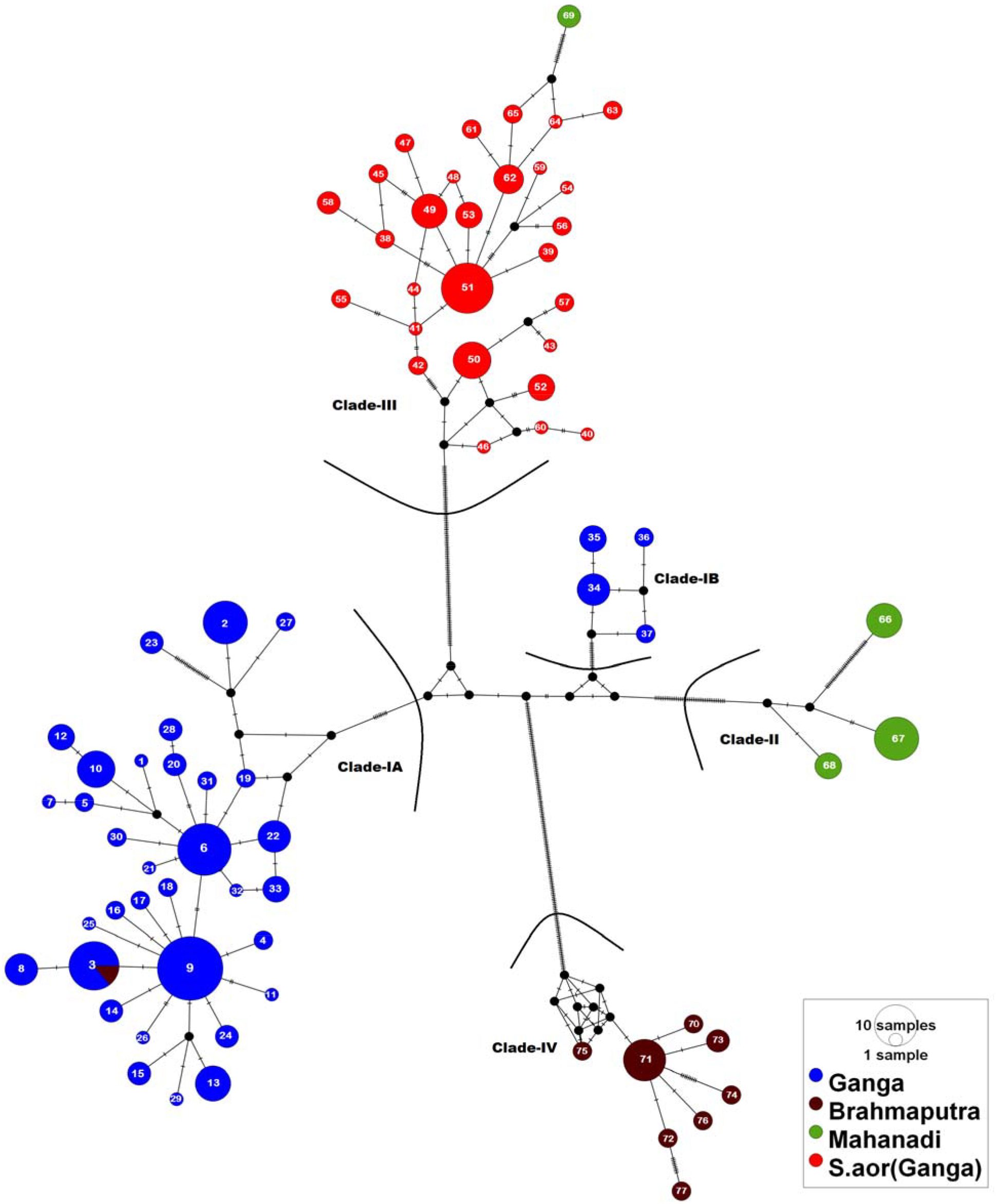
Median-joining network of *Sperata* species indicates the presence of three distinct clades. Clade-*I* consists of haplotypes of *S. seenghala* from the Ganges; Clade-*II* consists of Brahmaputra, Clade-*III* belongs to Mahanadi, and Clade-*IV* consists of haplotypes of *S. aor* from the Ganges.

### Genetic differentiation

A few sequences of Brahmaputra and Mahanadi were merged with Ganga *seenghala* and *S. aor*, respectively, therefore the factual mean pairwise genetic distance was estimated based on their clustering pattern. The analyses demonstrated significant genetic differentiation between all the lineages of *Sperata*. The genetic distinction between two known species, i.e, *S. seenghala* (Clade IA) and *S. aor* (Clade III) is 0.175. Considering these estimates as a reference for species-level variation, the Brahmaputra population is genetically highly diverse (0.25 to 0.28) than the other studied lineages. The genetic differentiation between two lineages of *S. seenghala* (Clade IA and IB) from Ganga is 0.048, whereas it was comparably high with Mahanadi 0.076 to 0.080 (Table. 3). The analysis of molecular variance indicated a high degree of structuring among the populations. A large proportion of genetic variation was attributable to the difference among the group, 62.56% and the differentiation among populations within the groups, 29.46% (Table 4). The *F*_*SC*_ and *Fst* values were found to be significant P<0.05.

**Table 3.**
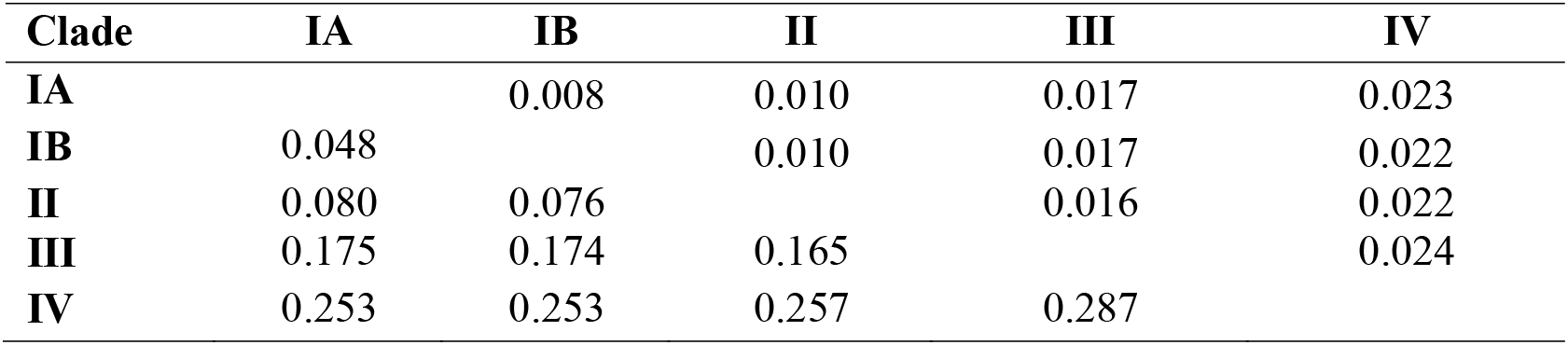
Estimates of evolutionary divergence over sequence pairs between groups of *Spereta* from four different rivers. Standard error estimate(s) are shown above the diagonal.

| Clade | IA | IB | II | III | IV |
| --- | --- | --- | --- | --- | --- |
| IA |  | 0.008 | 0.010 | 0.017 | 0.023 |
| IB | 0.048 |  | 0.010 | 0.017 | 0.022 |
| II | 0.080 | 0.076 |  | 0.016 | 0.022 |
| III | 0.175 | 0.174 | 0.165 |  | 0.024 |
| IV | 0.253 | 0.253 | 0.257 | 0.287 |  |

**Table 4.** Hierarchical analysis of molecular variance (AMOVA) for *S. Seenghala* from Ganges, Mahanadi, and Brahmaputra

| Source of variation | d.f. | Sum of squares | Variance components | % of variation | Fixation indices | P values |
| --- | --- | --- | --- | --- | --- | --- |
| Among groups | 3 | 11195.790 | 45.5 | 62.56 | FCT: 0.625 | 0.11±0.01 |
| Among populations within groups | 1 | 552.709 | 21.43 | 29.46 | FSC: 0.786 | 0.000 |
| Within populations | 279 | 1619.695 | 5.80 | 7.98 | FST: 0.920 | 0.000 |

### Demographic history

The demographical dynamics of the *Sperata* was inferred from the neutrality test and mismatch distributions (Table 2 and Fig. 4). No statistical significance values (P > 0.10) for Tajima’s D and Fu’s Fs were observed in *Sperata* populations. The bimodal shaped graphs were observed in *seenghala* of Ganga and Brahmaputra, whereas it was multimodal and ragged-shaped in *S.aor* and Mahanadi, suggested population subdivision and indications of demographic equilibrium. To assess the fitment of our data, we calculated the SSD and raggedness statistic (r-index) under the demographic expansion model for each population. However, these values were not statistically significant except Mahanadi, which indicates that neither the neutrality test nor the mismatch distribution test supported the hypothesis that the *Sperata* populations of the have passed through population expansions (Table 2).

**Figure 4.**
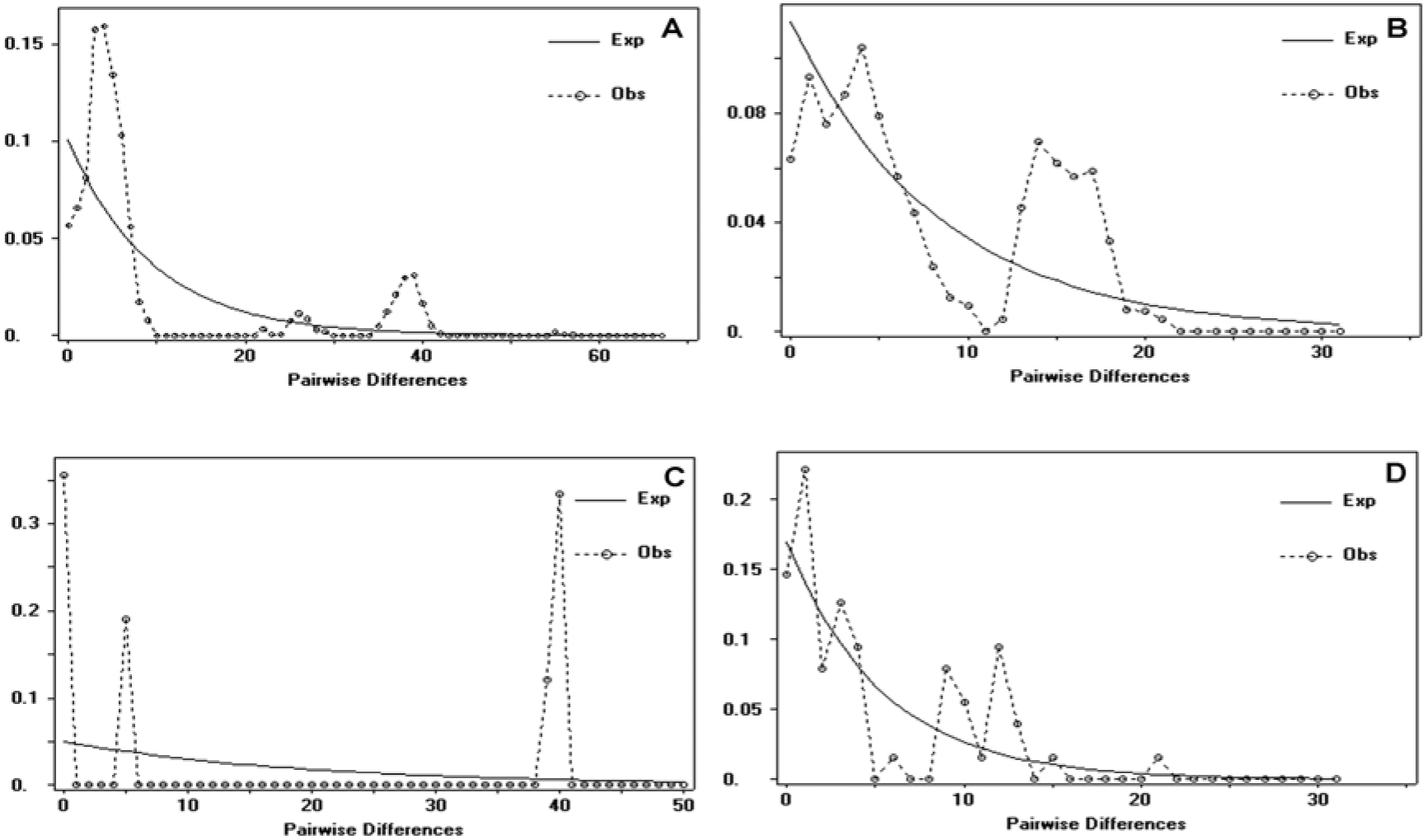
Mismatch distribution graph for *Sperata* populations A: Ganges *S. seenghala*; B: Ganges *S. aor*; C: Brahmaputra; and D: Mahanadi population. The x-axis shows the number of pairwise differences; the y-axis shows the frequency of pairwise comparisons. The observed and expected frequency under the population expansion model is represented by a dotted and continuous line.

## Discussion

The genetic structure and diversity analysis of any species that inhabits under high threats give us vital information for their development and effective management (Kumar *et al.*, 2017; Gupta *et al.*, 2018). For the conservation and to maintain the pure genetic stock, knowledge about the existing lineages is essential. Through this study, we first time generated the molecular data of *S. aor* and compared it with other existing populations of *Sperata spp*. Our data revealed a high haplotype diversity (>0.71) in *Sperata* species found in three Indian rivers. The pairwise genetic distance, phylogenetic, and AMOVA analysis indicated population structuring in *Sperata* spp. The phylogenetic relationship and network revealed the presence of two mtDNA lineages (clade IA and IB) of the *S. seenghala* from the Ganga river. Clade IA widely distributed throughout the Ganga, whereas Clade IB confined to lower stretch of Ganga, i.e, Munger district, Bihar. Interestingly, when we compared the data with Kumari *et al*. (2017), the two sequences (accession no. KT022220 and KT306656) from the Brahmaputra were merging with the Ganga haplotype (Hap.3). In contrast, three sequences (accession no. KT022199, KT022220, and KT022201) from Mahanadi (Hap. 69) were clustered with *S. aor* of the Ganga river. It indicated that the Brahmaputra River also sustains the gene pool of the Ganga river. The possible explanation for the existence of the Brahmaputra lineage in the Ganga is that both the rivers merge at the Sundarbans area of Bangladesh and provide the potential route for migration. Besides, the presence of similar gene-pool of Ganga S. *seenghala* in Brahmaputra river could be the effect of human-mediated propagation, and there is also a possibility of cross-contamination or cataloging error while doing genetic analysis. Further, three samples collected in a previous study from the Mahanadi river as *S. seenghala* belongs to *S. aor*. The clustering of Mahanadi *seenghala* with S. *aor* might be the result of the misidentification of species during sample collection. Therefore, the genetic differentiation was estimated between the different identified lineages observed in the study. Considering two known species

*S. seenghala* and *S. aor* as a reference, we observed high genetic differentiation value (17.5%) than the newly identified lineage of *seenghala* from Ganga (4.8%) and Mahanadi (8.0%). Interestingly, the genetic differentiation between Ganga-Brahmaputra was much higher (25.3%) than the values observed from *S. seenghala* and *S. aor*. This result indicated the presence of highly diverse lineages in the Brahmaputra than the *S. seenghala*, that needs to be confirmed based on their detailed morphological characteristics. The existence of structuring in *Sperata* is supported with a previous RAPD based study where population sub-structuring was observed in giant river catfish of Sutlej and Beas from the Indus river system (Saini, Dua & Mohindra, 2010). The study on stock discrimination using microsatellite markers from the middle to lower stretch of river Ganga, i.e, Narora–Kanpur, Varanasi, and Bhagalpur, revealed three different stocks of *S. aor* (Nazir and Khan, 2017). However, this result was contradictory with our mtDNA based study where we did not found any significant barrier to gene flow and structuring in *S.aor* across the sampling sites of the Ganga river.

Moreover, the existence of two distinct lineages of *S. seenghala*, which is diverse than *S.aor* and Mahanadi lineage, indicated the presence of multiple clades of *Sperata*. In the stretch of Varanasi to Sahebganj of the Ganga river, a large number of tributaries such as Gomti, Ghaghara, Gandak, Son, Kiul and Kosi merge with the Ganga that might have resulted in the migration and confinement of the different gene-pool of these rivers to a particular habitat (Nazir and Khan, 2017). Recently, Acharya *et al* (2019) reported population sub-structuring in the Brahmaputra, Ganga, Godavari, Mahanadi, and Narmada. The results indicated the low genetic divergence between Ganga–Brahmaputra than the Mahanadi populations, with comparatively low genetic diversity from the Ganga river due to the presence of excess homozygotes. In contrast to this, our mtDNA results showed Ganga *seenghala* is genetically close to Mahanadi than the Brahmaputra, which is supported by a previous study based on cytochrome *b* and D-loop region (Kumari *et al*. 2017). Though, incongruence results obtained from mtDNA and microsatellite analysis, suggesting that sex-biased dispersal behaviors that could contributed to disparities in genetic structuring of species. It also appears that the Ganga and Mahanadi are geographically more isolated than Brahmaputra and all three rivers obviously on an independent evolutionary trajectory; however, our study indicated that Brahmaputra lineages qualify the species level variation and adequately address the highly diverse lineages among S*pereta*. The non-significant Tajima’s D and Fu’s Fs tests suggested that there was no historical reduction in effective population size in these rivers. The mismatch distribution analysis showed bi-modal and multimodal mismatch distribution indicates the demographic stability or results of population admixture in S*pereta* that could be due to their reliability with allopatric divergence (Rogers & Harpending, 1992; Zhao et al. 2008).

The present genetic features in Ganga as well as other existing *Spereta* populations are the consequence of long-term geographical isolation and adaptation to the local environment.

## Conclusions

The present study provides the reference database for the identification and genetic differentiation of *S. aor* and *S. seenghala* from the Ganga river. The study highlighted the presence of two distinct lineages of *S. seenghala* from the Ganga river. Also, it indicated that the Brahmaputra population is much diverse than *S. seenghala* and *S. aor*. Therefore, the present study highlighted that the Brahmaputra giant river catfish is a highly diverse lineage that genetically qualifies the status of distinct species. However, it requires a detailed morphological study from more locations for the identification of multiple lineages existing in the Indian river that can formulate the appropriate conservation and management plans. This study indicated the presence of multiple lineages of *Spereta* from Ganga, Mahanadi, and Brahmaputra that considered as a distinct Evolutionary Significant Unit (ESUs). These findings will help understand the fish biology and implement the proper conservation, management plan, for development catfishes. However, detailed comparative morphological evidence with support of molecular data is required to be able to support their consideration as separate species.

## Acknowledgments

This study was supported by the Ministry of Jal Shakti. We acknowledge the support provided by the Director and Dean, WII, and the entire project team of NMCG. We thank state forest departments of Uttar Pradesh, Bihar, and Jharkhand for their support.

## Additional Information

### Author Contributions

SAH developed the project and acquired funds and permission for conducting the study. PY, AK, SAH and SKG conceptualized the methodological framework of the genetic and designed experiment. PY and AK collected the biological samples. PY, AK, NY, and MB performed wet lab work and prepared maps. PY, AK, and SKG performed the statistical analysis and wrote the manuscript. All authors approved the final version of the paper.

